# Experimental evidence for the gaze-signaling hypothesis: White sclera enhances the visibility of eye-gaze direction in humans and chimpanzees

**DOI:** 10.1101/2021.09.21.461201

**Authors:** Fumihiro Kano, Yuri Kawaguchi, Hanling Yeow

## Abstract

Hallmark social activities of humans, such as cooperation and cultural learning, involve eye-gaze signaling through joint attentional interaction and ostensive communication. The gaze-signaling and related cooperative-eye hypotheses posit that humans evolved unique external eye morphology, including exposed white sclera (the white of the eye), to enhance the visibility of eye-gaze for conspecifics. However, experimental evidence is still lacking. This study tested the ability of human and chimpanzee participants to detect the eye-gaze directions of human and chimpanzee images in computerized tasks. We varied the level of brightness and size in the stimulus images to examine the robustness of the eye-gaze directional signal against visually challenging conditions. We found that both humans and chimpanzees detected gaze directions of the human eye better than that of the chimpanzee eye, particularly when eye stimuli were darker and smaller. Also, participants of both species detected gaze direction of the chimpanzee eye better when its color was inverted compared to when its color was normal; namely, when the chimpanzee eye has artificial white sclera. White sclera thus enhances the visibility of eye-gaze direction even across species, particularly in visually challenging conditions. Our findings supported but also critically updated the central premises of the gaze-signaling hypothesis.

## Introduction

Humans employ eye-gaze in conspecific communication not only to infer others’ attentional foci, but also to check joint engagement in a cooperative task, and even to exchange information about own and others’ communicative intentions during key group activities such as cooperation and cultural learning [1, 2]. Closely related species such as nonhuman great apes also excel at employing own and others’ gaze in social interaction and various cognitive experiments [3–6]. However, previous studies have also reported several critical differences between human and nonhuman apes in their use and interpretation of own and others’ gaze in the context of gaze-following, joint engagement and ostensive communication [7–9]. One influential hypothesis, the gaze-signaling hypothesis [10, 11], later developed as the cooperative-eye hypothesis [8], proposes that humans have evolved special morphological features in the eye, including white (unpigmented) exposed sclera, to enhance the visibility of eye-gaze directions and thereby aid conspecifics to see one another’s visual targets without much attentional effort during social interaction. However, recent morphological studies have questioned this hypothesis based on new findings showing that humans are not necessarily unique among great apes in their external eye features [12–14], such as iris-sclera color contrast, eye shape, and the degree of sclera whiteness [also see, 15, 16, 17]. Nonetheless, those studies generally agree that one distinguishing feature of the human eye is uniform whiteness which is visible in its widely exposed sclera, a trait present in nearly all individuals. Currently, a key missing piece of information in the literature is experimental evidence answering the question of whether the human white sclera serves any communicative function for eye-gaze signaling.

To examine this question, we tested both humans and chimpanzees on their ability to distinguish the gaze direction of both chimpanzee and human faces via a computerized gaze-detection task using our unique set of eye stimuli (Figure 1). The task for human and chimpanzee participants was to distinguish gaze directions of human and chimpanzee eye images on a computer monitor (either in a keypress or visual search task; Figure 1B). Our experimental design was built based on previous experimental studies on gaze perception [8, 18–20]. To test our hypotheses (detailed below), we customized our design in four key aspects. First, our experiment uniquely employed a cross-species design, presenting the stimuli of both species to participants of both species. Second, we trained chimpanzees for a touch-panel task following a previous study [19] which successfully trained a chimpanzee to reliably differentiate eye-gaze directions of facial images. We did not use spontaneous gaze-following/cuing task mainly for a practical reason because previous studies have reported that chimpanzees do not often follow another’s eye-gaze direction, although they reliably follow another’s head direction [8, 21, 22]. Third, we uniquely tested the effect of visual noises, such as darkening and distancing, on the visibility of eye-gaze direction by manipulating the brightness and size of eye images because the strength of visual signal should critically depend on its degradation caused by natural noises [23]. Finally, inspired by a previous study with human participants [20], our stimuli included eye images in both normal and inverted colors (Figure 1). Specifically, as the human eye has white sclera and a darker iris, while the chimpanzee eye has dark sclera and a brighter iris, we simply inverted the lightness component of eye color in each stimulus species to make artificial dark sclera in the human eye and white sclera in the chimpanzee eye without affecting the iris-sclera color contrast (see Figure S1). This manipulation was made based on recent morphological studies showing that iris-sclera color contrast does not differ between humans and chimpanzees [12, 13, but see 14].

**Figure 1.**
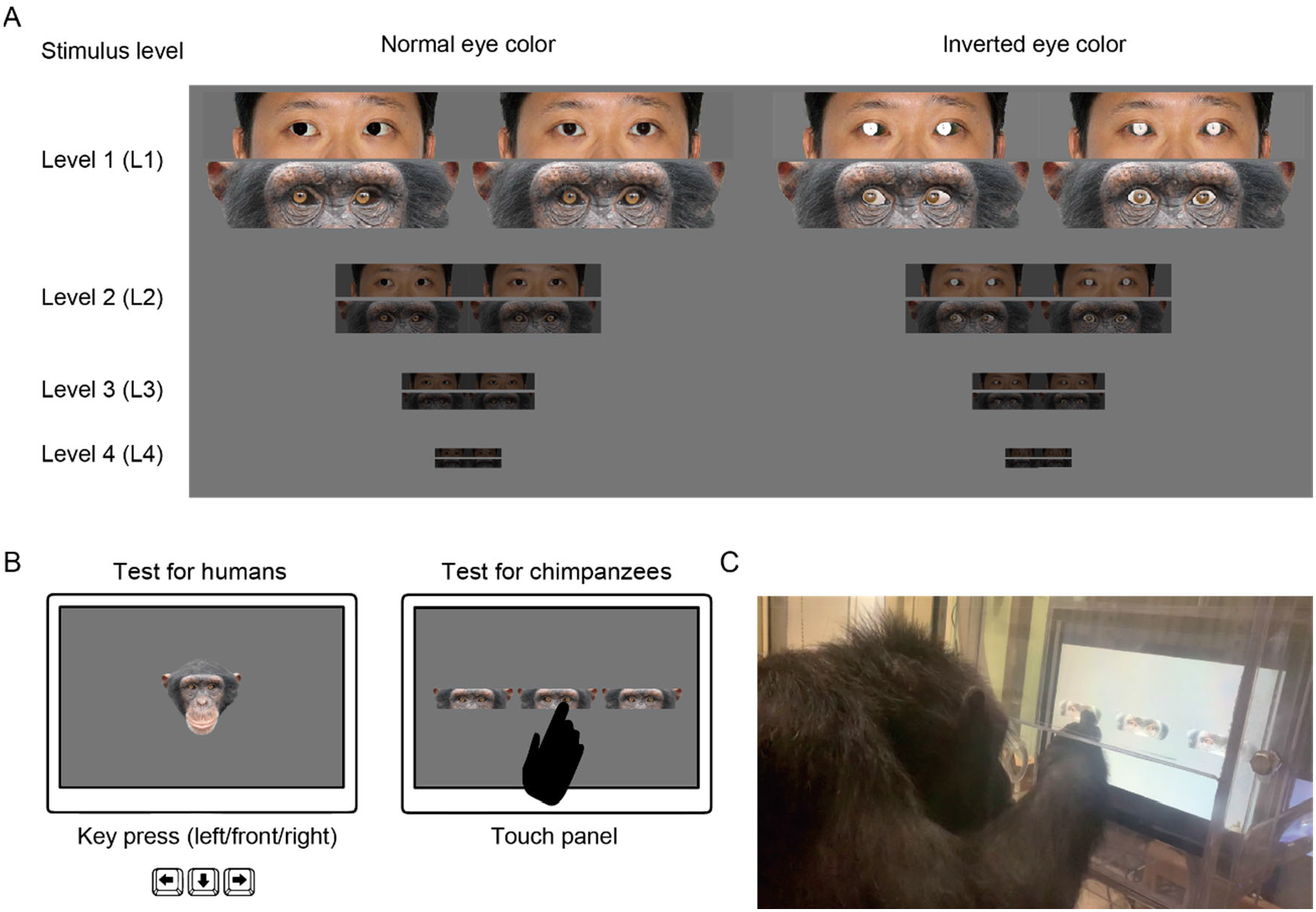
A. Experimental stimuli used in this study. Stimuli consisted of chimpanzee and human eye images with direct and averted (20 degrees) gaze in normal and inverted eye color at distinct levels of stimulus size and brightness. Permission was obtained to publish the human image (this image was only for presentation purposes; not used in this study but edited following the methods used in this study; see [24] for the stimuli used in the study). B. Schematics for the tests with human and chimpanzee participants. Participants of both species were presented with the stimuli of both species. In each trial, human participants indicated the gaze direction of stimulus face by key press (left/front/right), and chimpanzee participants indicated the averted gaze face among the two direct gaze faces by a touch response. C. Experimental setup for chimpanzees.

We developed 5 sets of hypotheses and predictions based on previous findings suggesting possible perceptual advantages of eye features and perceptual expertise of participants (Table 1). H1 posits the perceptual advantage of iris-sclera color contrast [12–14] and thus predicted no performance difference between the stimulus types in participants of both species. H2 is our key hypothesis which posits a perceptual advantage of white sclera [10, 11] and thus predicted increased performance in trials presenting the human eye in normal color and the chimpanzee eye in inverted color for participants of both species. H3 posits a perceptual advantage of the horizontally elongated shape, another distinguished feature of the human eye [10, 11, 13, 15, 25] (also see Figure S1) and thus predicted increased performance in trials presenting the human eye in both normal and inverted colors. H4 posits perceptual expertise for own-species eye colors in participants and thus predicted increased performance in trials presenting the images with own-species eye colors, namely white sclera and dark iris for human participants, and dark sclera and bright iris for chimpanzee participants. Such perceptual expertise has been demonstrated previously in experiments with human participants [20]. H5 posits perceptual expertise for normal eye colors in participants and thus predicted increased performance in trials presenting the eye (of either species) in normal color. H5 is built on a general possibility that gaze sensitivity may develop in individuals through repeated exposures to others’ gaze (in normal color) during daily interaction.

**Table 1.**
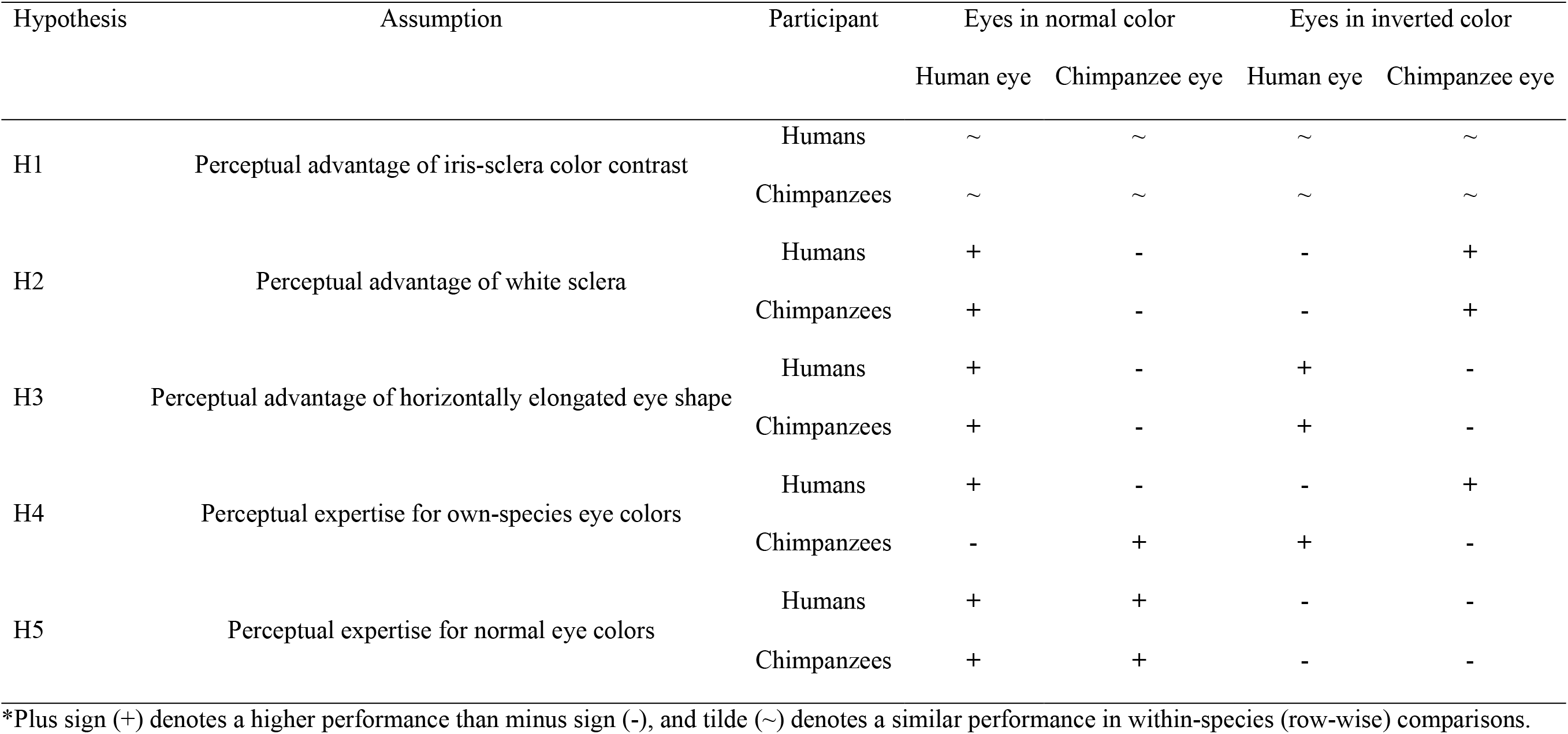
Hypotheses predicting the effect of eye color and shape on the visibility of eye-gaze in chimpanzee and human participants.

## Result

In our first study (Study 1), we tested 25 adult human participants (14 females, 11 males) in two experiments. All participants had moderate to extensive experience interacting with chimpanzees which ensured that they were familiar with the natural eye color patterns of chimpanzees and had experience reading their gaze direction. Experiment 1 presented participants with the stimuli of both humans and chimpanzees in normal colors at 4 stimulus levels (L1-4) varying in size and brightness. We tested the effect of stimulus level and species on participants’ accuracy in Generalized Linear Mixed Model (GLMM) and found that human participants performed better in trials presenting the human stimuli than those presenting the chimpanzee stimuli (χ^2^ = 37.08, d.f. = 1, P < 10^−8^), while their performance was worse in trials presenting smaller and darker stimuli (χ^2^ = 45.85, d.f. = 3, P < 10^−9^; see Table S1 for the full GLMM results) (Figure 2A). Experiment 2 presented the same participants with the stimuli in both normal and inverted colors at stimulus levels L3 and L4. We tested the effect of stimulus level, species, and color on participants’ accuracy in GLMM and found significant three-way interaction effects between these factors (Figure 2B; χ^2^ = 17.57, d.f. = 1, P < 10^−4^; also see Table S1). We then performed simple effects tests to examine the observed interaction effect further (Table S1 and Figure 2B). Critically, we found that inverting the color of the chimpanzee eye (with white sclera and a darker iris) significantly increased participants’ performance, while inverting the color of the human eye (dark sclera and a brighter iris) significantly decreased participants’ performance.

**Figure 2.**
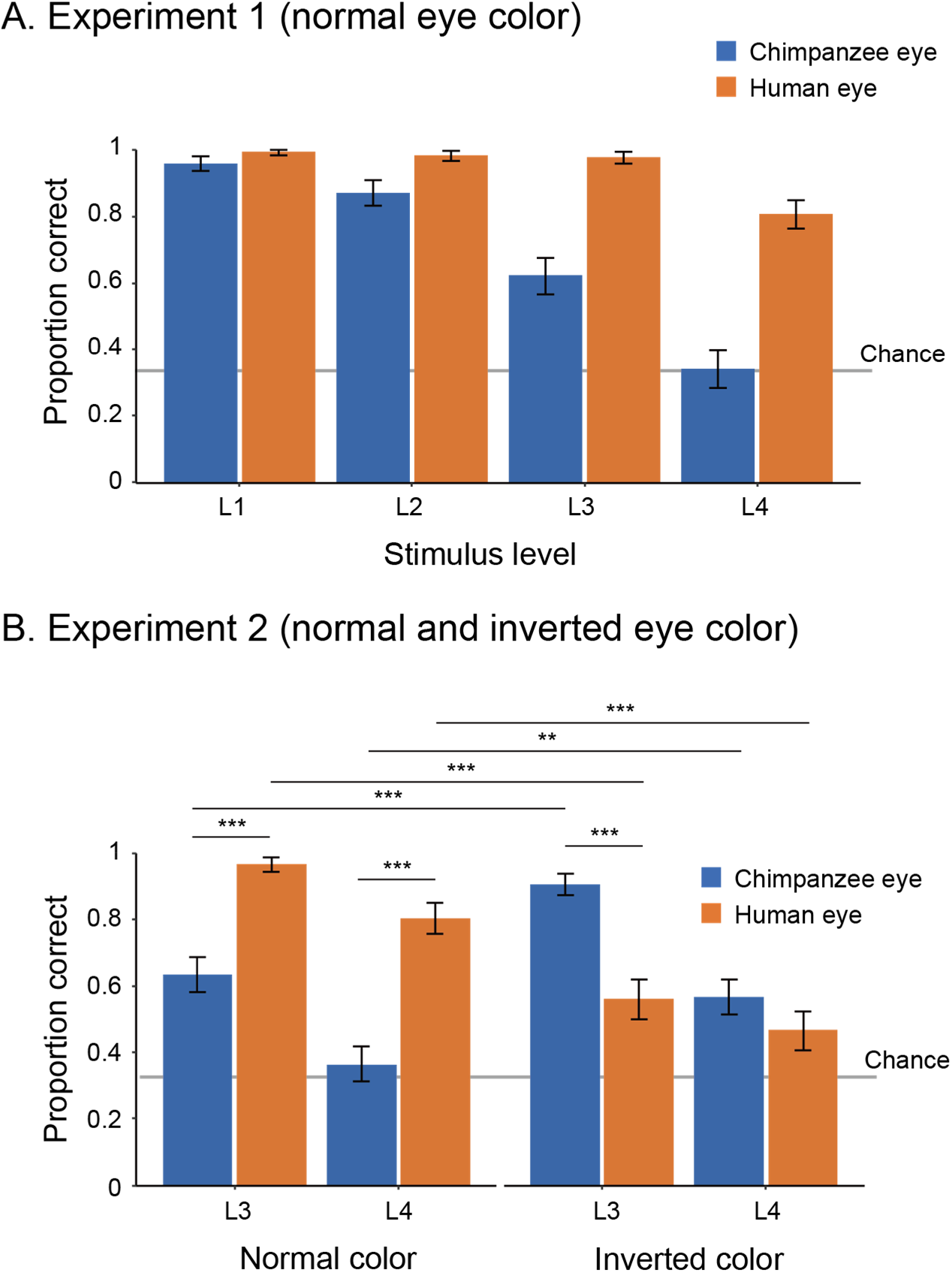
Performance of human participants in Study 1, represented as mean proportion correct in Experiment 1 (A) and 2 (B). Error bars are 95% confidence intervals based on a nonparametric bootstrap. Asterisks indicate significance in the post-hoc models ran for Experiment 2 (***p < 0.001, **p < 0.01).

In our second study (Study 2), we developed a task for chimpanzees to assess their ability to distinguish eye-gaze directions based on a previous study [19]. As in this previous study, our methods required extensive training for chimpanzees to reliably distinguish eye-gaze directions of stimuli (see Table S3 for details about training stages). Therefore, while we began training with 10 chimpanzees, only three (Natsuki, Hatsuka, Pendesa, all females, see Table S2 for further details) passed all the required training and test phases (see Table S4 and Figure S2). Study 2 involved two experiments. Experiment 1 tested those three chimpanzees and presented them with the human and chimpanzee eye images in normal color. To test chimpanzees at stimulus levels higher than L1 (i.e. darker and smaller), we gradually incremented the stimulus level (by 0.5) across sessions when individuals showed high performance in target trials presenting a given stimulus level (above 85% in 2 successive sessions). The test phase was defined as the sessions presenting stimulus level higher than (or equal to) L2.5 given that we observed clear performance differences between stimulus species at stimulus levels higher than L2 in Study 1 (see Table S5 for the number of sessions that each individual had in the pre-test and test phases). We tested chimpanzees’ accuracy during the test phase across repeated sessions at the individual level (with the alpha level corrected for the number of individuals in the Bonferroni correction, *α* = 0.05/3). Each chimpanzee completed a minimum of 20 test sessions. To avoid the ceiling effect, we incremented the stimulus level also during the test phase when chimpanzees showed high performance in target trials based on the same criteria. We tested the effect of stimulus species in GLMM on each chimpanzee’s accuracy during the test phase and found that all chimpanzees performed significantly better for the human stimuli than the chimpanzee stimuli (Natsuki: χ^2^ = 8.28, d.f. = 1, P = 0.004; Hatsuka: χ^2^ = 9.50, d.f. = 1, P = 0.002; Pendesa: χ^2^ = 21.94, d.f. = 1, P < 10^−5^; also see Table S1).

Experiment 2 tested two (Natsuki and Hatsuka) out of the three chimpanzees. Pendesa was dropped from this experiment because she took about twice as many training sessions as the other two chimpanzees (Table S4 and Figure S2). Experiment 2 presented them with the eye stimuli in inverted color in the first test phase (Test-B) and then the eye stimuli in normal color in the next test phase (Test-A2); thus, together with the results from Experiment 1 (also called Test-A1 phase), we tested chimpanzees in the ABA design. The Test-A2 phase started from stimulus level L3, which these two chimpanzees reached during the Test-A1 phase. The other procedures were identical with Experiment 1 (with *α* = 0.05/2). We compared each chimpanzee’s accuracy in target trials across Test-A1 and Test-B in GLMM and found a significant interaction effect between stimulus species and phase in both chimpanzees (Natsuki: χ^2^ = 34.61, d.f. = 1, P < 10^−8^; Hatsuka; χ^2^ = 8.39, d.f. = 1, P = 0.004; also see Table S1). We then compared each chimpanzee’s performance across Test-B and Test-A2 and found a significant interaction effect between the two factors in both chimpanzees (Natsuki: χ^2^ = 37.04, d.f. = 1, P < 10^−8^; Hatsuka; χ^2^ = 33.75, d.f. = 1, P < 10^−8^). To examine these observed interaction effects further, we performed simple effects tests (Table S1 and Figure 3). Critically, we found that both chimpanzees’ performance significantly increased from Test-A1 (normal color) to Test-B (inverted color) and then their performance decreased from Test-B to Test-A2 (normal color) in trials presenting the chimpanzee stimuli. Natsuki’s performance significantly decreased from Test-A1 to Test-B and then increased from Test-B to Test-A2 in trials presenting the human stimuli. Hatsuka’s performance did not significantly decrease from Test-A1 to Test-B but significantly increased from Test-B to Test-A2 in those trials.

**Figure 3.**
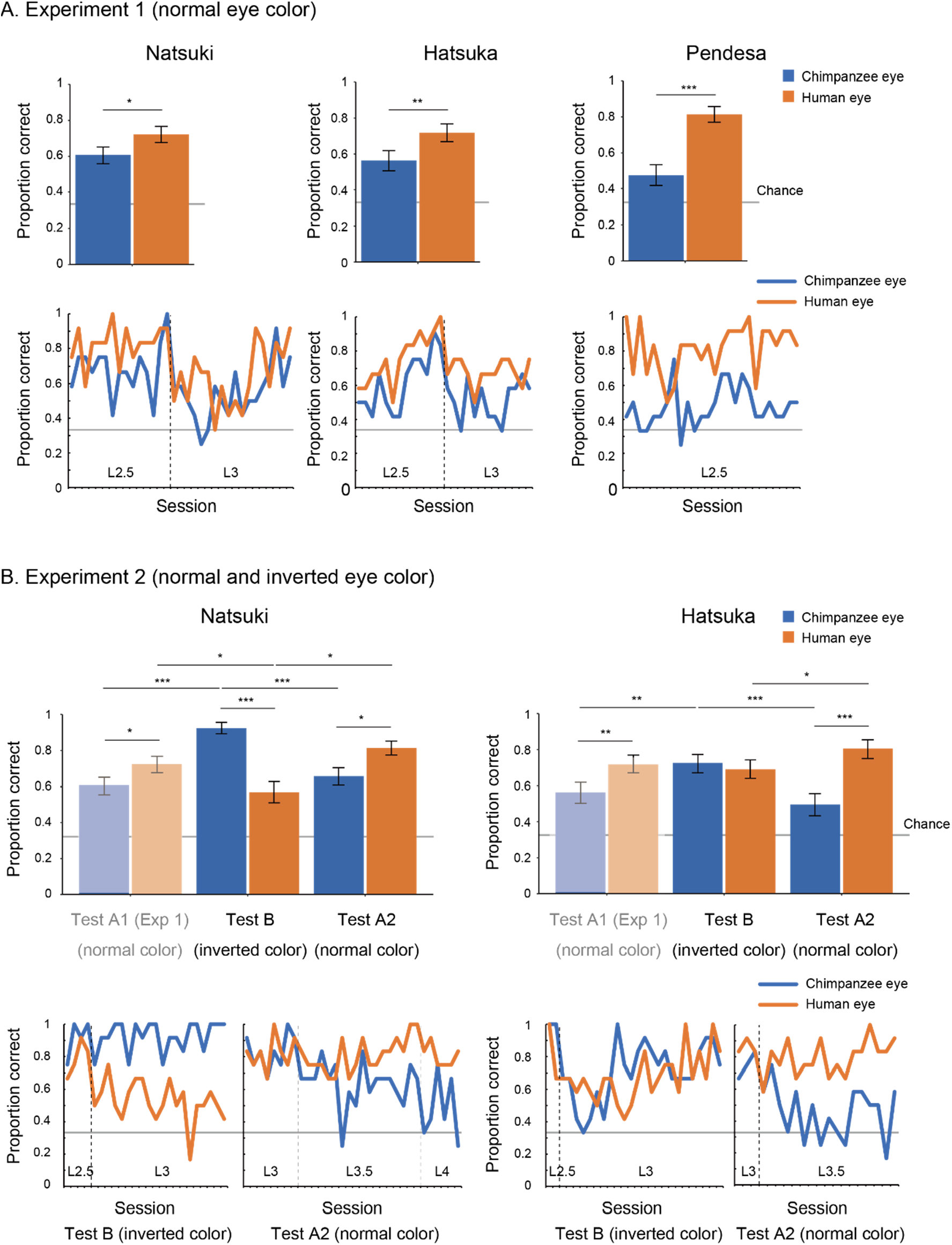
Performance of chimpanzee participants (Natsuki, Hatsuka, and Pendesa) in Study 2, represented as mean proportion correct calculated from all sessions (bar graphs) and each session (line graphs). Participants were tested in the ABA design. Specifically, Experiment 1 (A) presented the eye images in normal color (also termed Test-A1), and Experiment 2 (B) presented the eye images in inverted color (Test-B) and then those in normal color (Test-A2). The bar graphs of Experiment 2 repeat the same bar graphs of Experiment 1 (with their colors toned down) to aid comparisons in the ABA design. Dotted lines in line graphs denote the increment of stimulus level. Error bars in bar graphs are 95% confidence intervals based on a nonparametric bootstrap. Asterisks indicate significance in Experiment 1 and the post-hoc models ran for Experiment 2 (***p < 0.001, **p < 0.01, *p < 0.05; these significance levels were corrected for the number of individuals in each experiment).

## Discussion

Overall, these results revealed a striking perceptual advantage of white sclera for both human and chimpanzee participants for their ability to detect the eye-gaze direction of conspecific and allospecific images. Our results thus supported H2 (perceptual advantage of white sclera). Specifically, we found that white sclera, either in the normally-colored human eye or in the invertedly-colored chimpanzee eye, enhanced the visibility of eye-gaze in both human and chimpanzee images. We also found that, although both human and chimpanzee eyes are reliably visible when those eye stimuli were sufficiently large and bright (i.e. our L1 stimuli), the human eye-gaze was more visible than the chimpanzee eye-gaze particularly when those eye stimuli were in visually challenging conditions. H1 (perceptual advantage of iris-sclera color contrast) does not explain this result likely because it supposes only iris-sclera color contrast but not eye outline [10]; the latter is generally more conspicuous and likely more robust to darkening and distancing in the eye with white sclera than that with dark sclera. Our other hypotheses (H3-5) can explain our results partially, for either participant species or either stimulus species, but does not explain overall pattern of our results in a cross-species design. It should be noted that our results should not be interpreted as dismissing the assumptions of these other hypotheses but specifying the one which holds even in this cross-species design (i.e. H2). Nonetheless, H2 did not explain some of our other results; specifically, human participants (collectively) performed similarly in trials presenting the human and chimpanzee stimuli in the L4-inverted condition, and also chimpanzee Hatsuka performed similarly in trials presenting the human and chimpanzee stimuli during the Test-B (inverted color) phase. These results can be explained by either or the combination of H1 and H3-5. Overall, however, our findings clearly supported H2, but critically added that the perceptual advantage of white sclera is most clearly observable in visually challenging conditions. This indicates that the key function of white sclera is to equip eye-gaze signal with robustness against its degradation caused by natural noises.

Consistency of our results across human and chimpanzee participants suggest that the visibility of human eye-gaze can be attributed to simple color properties of eyes and thus may be supported by basic perceptual mechanism shared among human and chimpanzee participants. Moreover, our study also showed that chimpanzees required extensive training to distinguish between different gaze directions of human and chimpanzee eyes, and many of our chimpanzees were unable to pass all the required training phases. This suggests that chimpanzees are not skilled at or motivated for distinguishing the eye-gaze directions of others, consistent with a previous gaze-following/cuing study showing that they generally rely on a more global cue (i.e. head orientation) to follow another’s gaze [8, 21, 22]. On the other hand, human infant follow another’s gaze [26] at very early ages and primarily rely on eye-gaze rather than head direction to do so [27]. Therefore, these results together suggest that, in line with the cooperative-eye hypothesis [8], humans may have co-evolved their sensitivity to others’ eye-gaze as well as their special eye feature to most effectively employ conspecific eye-gaze signals during social interaction.

In conclusion, we demonstrated that whiteness in exposed sclera enhances eye-gaze signaling. We thus provided experimental support for the gaze-signaling hypothesis despite recent criticisms on this hypothesis [12–14]. However, we also propose several significant updates on this hypothesis. Specifically, we found that it is white sclera but not necessarily other distinguishing features, such as iris-sclera color contrast and some variation in horizontal eye elongation, which critically distinguishes the human eye from the chimpanzee eye with respect to the visibility of eye-gaze direction. Moreover, we found that the key function of white sclera is to enhance the eye-gaze signal, so that it is more reliable, even in visually challenging conditions. These new findings, when combined with the original and related hypotheses, suggest that humans excel at distinguishing conspecific gaze direction, even across various challenging visual conditions (even in a shade or at some distance) and that this is a vital part of information acquisition and exchange for humans engaging in social and cooperative behaviors during their everyday group activities.

## Acknowledgments

We thank the staffs at Kumamoto Sanctuary and Primate Research Institute (especially, Etsuko Ichino, Ayumu Santa, Akiho Muramatsu, and André Gonçalves and Drs. Naruki Morimura and Ikuma Adachi) for assistance with performing experiments. We also thank Drs. Christopher Krupenye and Lydia Hopper for their edits and fruitful comments. Financial supports came from Japan Society for the Promotion of Science KAKENHI Grants 19H01772 and 20H05000 to F.K and 18J20077 to Y.K.

## Author Contributions

F.K. designed the study; F.K., Y.K., H.Y. performed the experiments; F.K. drafted and all authors refined the manuscript.

## Methods

Participants. Study 1 tested 25 human adults (14 females, 11 males; 24 East/South Asians and 1 Caucasian male) who had moderate to extensive experience in caretaking or studying chimpanzees (3 months = 1; 1-5 years = 10; 5-10 years = 4; > 10- years = 10). Our participants included 10 participants who had extensive experience interacting with chimpanzees (over a decade). We obtained the same results even when we restricted these analyses to those participants. All human participants were workers or students at Kumamoto Sanctuary (KS) or Primate Research Institute (PRI) who were directly invited to participate in this experiment. All were naïve to the experimental hypotheses in this study. All reported to have normal to corrected-to-normal vision and no color blindness. Written informed consent was obtained from all participants prior to the study. Experimental protocol was approved by the internal ethical committee for human experiments in PRI (No. 2020-05).

Study 2 trained 10 chimpanzees (9 females, 1 male). Among them, 3 chimpanzees (Natsuki, Hatsuka, Pendesa, all females) passed all the training stages and participated in Experiment 1. Two of these 3 chimpanzees (Natsuki, Hatsuka) participated in Experiment 2. Daily veterinary checks indicated no vision problems (including color blindness) in these chimpanzees that may practically interfere with the execution of current experiments. Chimpanzees lived in a social group of conspecifics at KS or PRI. All chimpanzees were tested in a dedicated testing room at each facility, and their daily participation was voluntary, in that they could decide whether to enter the testing room on a given testing day. They received regular feedings, daily enrichment, and had *ad libitum* access to water. Animal husbandry complied with institutional guidelines (KS: Wildlife Research Center “Guide for the Animal Research Ethics”; PRI: 2002 version of “The Guidelines for the Care and Use of Laboratory Primates”), and research protocol was approved by the institutional research committee (KS: WRC-2020-KS008A/009A; PRI: 2020-193/209). See Table S2 for details about participants.

Apparatus. Study 1 tested human participants in a standard office setting either in KS or PRI. Two participants were tested remotely online given the COVID-19 situation at the time of experiment. They received the same task program online and performed the task on their own computer in a standard office. Although slight differences existed in experimental setups between these two and the other participants (detailed below), we confirmed that including or not including them in our analysis yielded the same results. Participants sat in front of a 23-inch monitor (52.7×29.6 cm, SE2416H, Dell, Round Rock, USA; for one online participant, 52.2×29.3 cm, 243V5QHABA/11, Phillips, Amsterdam, Netherland; for the other online participant, 50.9×28.6 cm xub2390hs-b3, Iiyama, Japan; all monitors were in 1920×1080 pixels and set at 100% brightness and 50% contrast). They placed their 2^nd^ to 4^th^ digit fingers on the left, down, and right keys of the standard keyboard connected to the computer. With this setup, the viewing distance was about 60-70 cm. Participants were told to sit in front of the monitor as they normally would and not to move their original head position throughout the experiments.

Study 2 tested chimpanzee participants in a test room equipped with touch panels (ET1790L-7CWB-1-ST-NPB-G, Touch Panel Systems, Yokohama, Japan, in KS; LCD-AD172F2-T, IO-DATA, Kanazawa, Japan, in PRI; both 34.5×26.0 cm; both 1280×1024 pixels; both 100% brightness and 50% contrast) were installed with its center 45cm from the floor. With this position, the eye level of the chimpanzees was roughly at the center of the monitor when they sat on the floor. The monitor was installed 15cm behind a wall made of transparent polycarbonate panels, and there was a rectangle hole sized 40×15 cm on the panel so that chimpanzees could view the stimuli through the panel while making a touch response by inserting their arm through the hole (Figure 1). With this setup, the viewing distance was about 30-40 cm. This visual distance for chimpanzee participants is shorter than that for human participants, and thus overall task difficulty should be lower for them, consistent with other procedural differences that we made to ease the task difficulty for chimpanzees.

Stimuli. We prepared chimpanzee and human facial images in normal and inverted eye color with different levels of size and brightness. To create the chimpanzee stimuli, we selected 10 high-resolution facial images of chimpanzees of both sexes from KS in image collections obtained from colleagues. The selection criteria were as follows; 1) the photographed individual oriented both head and eye directly to the camera, 2) all eye features of the individual were clearly visible, 3) no strong shade was visible on the face, and 4) no expression was shown in the face. The selected chimpanzee individuals included juveniles and adults of both sexes, and their sclera colors were uniformly dark (from the iris edge to the eye corner). To create the human stimuli, we selected 10 high-resolution facial images of humans from the image collections published for research use [24] based on the same criteria as above. The selected human individuals are teenagers of both sexes in various ethnicities with various skin and eye colors, and their sclera colors were uniformly white (from the iris edge to the eye corner). We balanced the selection of human individuals so that we could include a wide variety of skin and eye colors among the stimulus individuals in our experiments (and we controlled for a potential effect of such variations including stimulus individuals as a random factor in our mixed models; see below). Study 1 used the whole sets of chimpanzee and human stimuli, which consisted of 10 stimulus individuals in each stimulus species, and Study 2 used 6 stimulus individuals in each stimulus species (due to the procedural differences between studies; see below).

The facial images of both chimpanzees and humans were then cropped to include only face and hair and automatically level-adjusted to reduce the variations in overall brightness across images in Photoshop (Adobe, San Jose, USA). Those cropped images were then pasted into a uniform 50 %-gray background sized 400×400 pixels. The size of the cropped facial image (for both chimpanzee and human image) was adjusted based on its iris diameter, which was set at 16 pixels (4.2-4.4 mm on the monitors used in both Study 1 and 2) in all images (of both stimulus species). This size adjustment was performed to test the effect of eye shape (horizontal elongation of eye opening, related to H3) independently from its absolute size and also to test the effect of white sclera independently from its exposed area (see below for the quantification of these parameters). It should be noted that, due to these controls, our chimpanzee stimuli were presented as slightly larger than the size proportional to human stimuli because the eyeball size of human is slightly (about 5-10%) larger in that of chimpanzee [28–30]. To create the facial images with averted gaze, the eyeball part of each face (with direct gaze) was cropped and then shifted 6 pixels to the side (this corresponded to the rotation of the eyeball of about 20 degrees in both stimulus species). We then filled the blank areas in the shifted eye by copying the sclera colors of the original image using the “stamp” tool in Photoshop. To create the facial images with inverted eye color, we first cropped the eye openings of each face (with both direct and averted gaze) and then inverted the lightness component of the cropped part (L value of Lab color) while keeping its color component (a and b values of Lab color) unchanged in a custom-made Matlab program (Mathworks, Natick, USA).

We then evaluated the shape and color of the eyes in our images using the methods that we customized based on previous studies [10, 12–14, 25]. We first created the Region-Of-Interest (ROI) mask respectively for iris and sclera by tracing and filling the edge of each feature in Photoshop and a custom-made Matlab program. We then calculated the color of each ROI as the mean of Lab color in all pixels within that ROI. We then calculated the color difference between iris and sclera in each image as a Euclidean difference between the mean values of these ROIs. We confirmed that the iris-sclera color differences did not significantly differ between the human and chimpanzee eye in both normal and inverted color (Figure S1). We also measured the shape of the eyes in each image using the same ROIs. We confirmed that the human eye was horizontally longer than the chimpanzee eye, but the size of sclera ROI did not differ between the human and chimpanzee eye (Figure S1).

Finally, we converted the facial images to various levels of size and brightness. Study 1 used 4 stimulus levels (L1-4). L1 stimuli measured 400 pixels in width (original) and 100% brightness (original), L2 stimuli measured 200 pixels in width (1/2) and 50% brightness (1/2), L3 stimuli measured 100 pixels in width (1/4) and 33% brightness (1/3), and L4 stimuli measured 50 pixels in width (1/8) and 25% brightness (1/4). These size and brightness levels were determined based on pilot experiments with 2 human participants (who did not participate in Study 1) so that the gaze direction of L4 stimuli was recognizable to both participants at least in one of the stimulus species in either normal or inverted eye color. In Study 2, we prepared 3 additional stimulus levels, L1.5, L2.5, and L3.5, which were the intermediate between L1 and 2, L2 and 3, and L3 and 4, respectively, in terms of size and brightness (i.e. L1.5: 300 pixels in width, 75% brightness; L2.5: 150 pixels in width, 42% brightness; L3.5: 75 pixels in width, 29% brightness) so that chimpanzees could move to the next stimulus level without showing substantial drops in their performances (see details about the test procedures below). Study 1 and 2 used identical stimuli except that Study 1 presented the whole face in a 1:1 square image (e.g. 400 × 400 pixels), while Study 2 presented only the eye region in a 4:1 rectangle image (e.g. 400 × 100 pixels) to reduce attentional demands on chimpanzees.

Task procedures: We made the task procedures of Study 1 and 2 as similarly as possible, although several unavoidable differences existed due to the fact that chimpanzees required extensive training to master the gaze-detection task. In Study 1, the task for human participants was to indicate the direction of gaze (left/front/right) in the stimulus face presented at the center of the screen by key press in each trial. They were instructed to answer as accurately and quickly as possible.

Study 1 consisted of two experiments. All participants completed Experiment 1 and 2 in this order. Experiment 1 presented the stimuli with normal eye colors at L1-4 levels. Experiment 2 presented the stimuli with both normal and inverted eye colors at L3-4 levels. Prior to each experiment, they completed 20 practice trials presenting L1 stimuli (with normal and inverted colors respectively for Experiment 1 and 2). Each experiment consisted of a total of 96 trials with 8 blocks (12 trials within each block). In both experiments, each block presented the stimuli of the same species, and the 8 blocks alternately presented the chimpanzee and human stimuli. In Experiment 1, each block started with 3 consecutive trials presenting L1 stimuli of either species (with normal eye colors) and then increased the stimulus level every 3 trials; namely, 1^st^-3^rd^ trials, 4^th^-6^th^ trials, 7^th^-9^th^ trials, and 10^th^-12^th^ trials respectively presented L1, L2, L3, and L4 stimuli of either species. Thus, in Experiment 1, 12 trials presented stimuli of either species (chimpanzee, human) at each stimulus level (L1-4). In Experiment 2, each block (12 trials) started with 6 consecutive trials presenting L3 stimuli of either species, and then 6 consecutive trials presenting L4 stimuli of the same species. The first two blocks (1^st^ and 2^nd^ blocks) presented the stimuli of the two species with normal eye colors (each block presented one species), and then the next two blocks (3^rd^ and 4^th^ blocks) presented the stimuli of the two species with inverted eye colors. The 5^th^ and 6^th^ blocks and the 7^th^ and 8^th^ blocks again presented the stimuli of the two species with normal eye colors and then inverted eye colors. Thus, in Experiment 2, 12 trials presented stimuli of either species (chimpanzee, human) at each stimulus level (L3, L4) in either eye color (normal, inverted). Each participant completed all experiments in 25-30 minutes. All experiments were conducted in November 2020.

The number of times in which each gaze direction (left/front/right) was presented was balanced in each participant (i.e. each direction was presented in 32 trials per participant), and the number of times in which each stimulus individual was presented was also balanced both within each participant and across participants (i.e. each stimulus individual was presented on average 4.8 times per participant). The order of presenting the chimpanzee or human stimuli in the first block was counterbalanced across participants. The orders of gaze directions (left/front/right) and stimulus individuals were pseudorandomized so that the same gaze direction was not presented in more than 2 successive trials, and the same stimulus individual was not presented in any successive trials.

In Study 2, the task for the chimpanzee participants was to indicate the image with averted gaze (shifted to the right; called the target image) among two other images (called the distractor images) with direct gaze by a touch response in each trial (i.e. 3-item visual search task, following the task design by [19]). The three images were centered at 220, 640 (center), and 860 pixels horizontally and 512 pixels vertically on a 1280 ×1024 pixels monitor. Chimpanzees were given a sip of grape juice or a piece of apple (depending on the chimpanzees’ preference) when they answered correctly in each trial (the same amount of reward was given for each chimpanzee throughout the study). Prior to training, we performed a pilot experiment (200-600 trials for each chimpanzee) to decide the general task design, especially in terms of the number of distractors in each trial, the number of trials in each session, and the features of initial stimuli, so that the chimpanzees could gradually learn the task.

As in Study 1, Study 2 consisted of 2 experiments. Experiment 1 presented the stimuli with normal eye colors, and Experiment 2 presented the stimuli with inverted eye colors and then the stimuli with normal eye colors. Thus, those two experiments presented the eye stimuli in normal and inverted colors in the ABA design. Throughout Study 2 (both training and test), each session consisted of 48 trials and 4 blocks. Each block (12 trials) presented the stimuli of the same species, and the 4 blocks alternately presented the chimpanzee and human stimuli. Each chimpanzee performed 1-8 sessions per day depending on their motivation. Each session lasted about 10 minutes. Study 2 took about 8 months from August 2020 to March 2021 including both training and test periods.

Training performed prior to these two experiments in Study 2 consisted of 6 training stages, and chimpanzees were trained for the task in a step-by-step manner through these stages. Training stage 1 presented the target image with no iris with the distractor images with irises (in direct gaze). Training stages 2-4 presented the target image in which the iris was positioned in 38, 30, and 20 degrees (the final iris position in Training 4), following the training procedure employed by a previous study [19]. Training 1-4 used two stimulus individuals per stimulus species, and Training stage 5 and 6 added two new stimulus individuals per stimulus species in each stage.; thus, Training 5 and 6 respectively presented 4 and 6 stimulus individuals per stimulus species (the final stimulus set in Training 6). The criterion of passing each training stage was either scoring over 90% in one session or 80% in two consecutive sessions both in trials presenting the chimpanzee stimuli and those presenting the human stimuli. We trained chimpanzees in the same number of trials for the human and chimpanzee stimuli to avoid biasing their learning for either stimulus species. Eight chimpanzees learned this visual search task with the most basic stimulus set (Training 1) and continued the next training stages (Training 2-6). During the latter training stages, there was no consistent bias across individuals in their performances for the trials presenting the human and chimpanzee stimuli. However, some chimpanzees performed notably better in trials presenting the chimpanzee stimuli than those presenting the human stimuli (Cleo, Iroha, Mizuki), and some showed an opposite pattern (Pendesa), and others performed similarly in those trials (Figure S2). Three chimpanzees (Natsuki, Hatsuka, Pendesa) passed all training stages after extensive training (Table S4) and learned to reliably differentiate eye-gaze directions of both human and chimpanzee stimuli (L1, normal eye color; Figure S2). Three chimpanzees passed all the training stages and then participated in Experiment 1. See Table S4 for the number of sessions each chimpanzee had in each training stage.

Experiment 1 was divided into pre-test and test phases (pre-Test-A1 and Test-A1 phases). To test chimpanzees at stimulus levels higher than L1, we incremented the stimulus level by 0.5 when the chimpanzee scored above 85% in 2 successive sessions (in all trials at stimulus level L1 and in test trials at stimulus level higher than L1; see below for details about the test and baseline trials). The pre-test phase started from stimulus level L1 and the test phase started from stimulus level L2.5. We used the stimulus level higher than (or equal to) L2.5 for the test phase because the results from Study 1 (human participants) indicated that clear performance differences between the stimulus species emerged at stimulus levels higher than L2. To examine individuals’ performance across sessions, each individual completed a minimum of 20 test sessions. To avoid the ceiling effect, we also incremented the stimulus level by 0.5 when the individual scored above 85% in 2 successive sessions in test trials during the test phase. Two chimpanzees (Natsuki and Hatsuka) participated in Experiment 2. The other chimpanzee (Pendesa) took more than twice as many sessions as the other two chimpanzees for training and thus was dropped from Experiment 2 (Table S4 and Figure S2). Experiment 2 first presented the stimuli with inverted eye colors (pre-Test-B and Test-B phases), and then presented the stimuli with normal eye colors (pre-Test-A2 and Test-A2 phases). As in Experiment 1, the pre-Test-B and pre-Test-A2 presented L1-2 stimuli and the Test-B and Test-A2 phase presented L2.5-4 stimuli. Each chimpanzee completed a minimum of 20 test sessions respectively in the Test-B and Test-A2 phase. The other procedures were identical to those in Experiment 1. It should be noted that, although the number of these test sessions varied across individuals and test phases, we confirmed that limiting the data set to 20 sessions in all individuals and sessions yielded the same results. See Table S5 for the number of sessions each chimpanzee had in each pre-test and test stages.

In both Experiment 1 and 2, (pre-)test sessions presenting the stimulus level higher than or equal to L1.5 consisted of 24 baseline and 24 test trials. The baseline trials presented L1 stimuli, and the test trials presented the stimuli at higher levels. Each block (12 trials) presented 6 baseline trials consecutively and then 6 test trials. Thus, in each test session, 12 trials presented stimuli of either species (chimpanzee, human) at the L1 (baseline trials) or higher levels (test trials). Chimpanzees kept high performances in baseline trials across sessions for both human and chimpanzee stimuli in the Test-A1 phase (L1 stimuli with normal color; Natsuki: 89% ± 12 vs. 90% ± 8; Hatsuka: 91% ± 10 vs. 89% ± 8; Pendesa; 97% ± 5 vs. 91% ± 7; mean ± SD), Test-B phase (L1 stimuli with inverted color; Natsuki: 91% ± 7 vs. 94% ± 7; Hatsuka: 92% ± 10 vs. 96% ± 6; mean ± SD), and Test-A2 phase (L1 stimuli with normal color; Natsuki: 91% ± 10 vs. 94% ± 7; Hatsuka: 92% ± 7 vs. 96% ± 5; mean ± SD).

As in Study 1, the number of times in which the target image (with averted gaze) was presented on each location (left/center/right) was balanced in each session (i.e. each gaze direction was presented in 16 trials per session), and the number of times in which each stimulus individual was presented was balanced in each session (i.e. each stimulus individual was presented 4 times per session). The order of presenting chimpanzee or human images in the first block was counterbalanced across sessions. The locations of the target images and the order of presenting the stimulus individuals were pseudorandomized so that the target image did not appear on the same location in more than 2 successive trials, and the same stimulus individual was not presented in any successive trials. See Table S3 for the summarized descriptions of each stage at the training and (pre-)test phases and Table S4 and S5 for the number of sessions in each stage at the training and (pre-)test phases, respectively.

Data analysis: To test the participants’ performance differences between conditions, we ran binomial GLMM (Generalized Linear Mixed Model) in R (version 4.0.5). In Experiment 1 of Study 1, the model included stimulus species (chimpanzee image, human image) and stimulus level (L1-4) as test fixed factors, the interaction between those test factors, block and (within-block) trial, which was nested in each block, as control fixed factors, and participant and stimulus individual as random factors. In Experiment 2 of Study 1, we used the same model with stimulus color (normal, inverted) as an additional test fixed factor (and its interaction with the other test factors). In Study 2 (chimpanzees), as we evaluated chimpanzees’ performance in repeated sessions and adjusted it by incrementing stimulus levels according to their performance, we treated session as a random factor and did not include stimulus level in the model. Therefore, the model included stimulus species as a test fixed factor, block and (within-block) trial as control fixed factors, and stimulus individual and sequence as random factors. Study 2 performed statistical tests for each chimpanzee with the alpha level adjusted for the number of individuals in the Bonferroni correction; namely, 0.05/3 in Experiment 1 and 0.05/2 in Experiment 2. For all models in Study 1 and 2, we included all possible random slope components, although we removed the correlations between random slopes and intercepts to handle the non-convergence issues [31]. Overdispersion was checked using the dispersion parameters which derived from the R package “blmeco” and did not seem to be an issue in any of our model (they ranged between 0.77 and 1.12). The significance of a given term was tested using a likelihood ratio test. Nonsignificant interaction terms were dropped to test the significance of lower-order terms. When the interaction term was significant, post-hoc comparisons were performed to examine simple effects at each factor level.

## References

1. Csibra, G., and Gergely, G. (2009). Natural pedagogy. Trends Cogn. Sci. 13, 148–153.

2. Tomasello, M., Carpenter, M., Call, J., Behne, T., and Moll, H. (2005). Understanding and sharing intentions: The origins of cultural cognition. Behav. Brain Sci. 28, 675–691.

3. Gomez, J.C. (1996). Ostensive behavior in great apes: The role of eye contact. In Reaching into thought: The minds of the great apes, A.E. Russon, K.A. Bard and S.T. Parker, eds. (New York: Cambridge University Press), pp. 131–151.

4. Okamoto-Barth, S., Call, J., and Tomasello, M. (2007). Great apes’ understanding of other individuals’ line of sight. Psychol. Sci. 18, 462–468.

5. Bräuer, J., Call, J., and Tomasello, M. (2005). All great ape species follow gaze to distant locations and around barriers. J. Comp. Psychol. 119, 145–154.

6. Kano, F., and Call, J. (2014). Cross-species variation of gaze following and conspecific preference among great apes, human infants and adults. Anim. Behav. 91, 137–150.

7. Carpenter, M., and Tomasello, M. (1995). Joint attention and imitative learning in children, chimpanzees, and enculturated chimpanzees. Soc. Dev. 4, 217–237.

8. Tomasello, M., Hare, B., Lehmann, H., and Call, J. (2007). Reliance on head versus eyes in the gaze following of great apes and human infants: The cooperative eye hypothesis. J. Hum. Evol. 52, 314–320.

9. Kano, F., Moore, R., Krupenye, C., Hirata, S., Tomonaga, M., and Call, J. (2018). Human ostensive signals do not enhance gaze following in chimpanzees, but do enhance object-oriented attention. Anim. Cogn. 21, 715–728.

10. Kobayashi, H., and Kohshima, S. (2001). Unique morphology of the human eye and its adaptive meaning: Comparative studies on external morphology of the primate eye. J. Hum. Evol. 40, 419–435.

11. Kobayashi, H., and Kohshima, S. (1997). Unique morphology of the human eye. Nature 387, 767–768.

12. Perea-García, J.O., Kret, M.E., Monteiro, A., and Hobaiter, C. (2019). Scleral pigmentation leads to conspicuous, not cryptic, eye morphology in chimpanzees. Proc. Nat. Acad. Sci. 116, 19248–19250.

13. Caspar, K.R., Biggemann, M., Geissmann, T., and Begall, S. (2021). Ocular pigmentation in humans, great apes, and gibbons is not suggestive of communicative functions. Scientific Reports 11, 12994.

14. Mearing, A.S., and Koops, K. (2021). Quantifying gaze conspicuousness: Are humans distinct from chimpanzees and bonobos? J. Hum. Evol. 157, 103043.

15. Mayhew, J.A., and Gómez, J.-C. (2015). Gorillas with white sclera: A naturally occurring variation in a morphological trait linked to social cognitive functions. Am. J. Primatol. 77, 869–877.

16. Perea-García, J.O., Danel, D.P., and Monteiro, A. (2021). Diversity in primate external eye morphology: Previously undescribed traits and their potential adaptive value. Symmetry 13.

17. Mearing, A.S., Burkart, J.M., Dunn, J., Street, S.E., and Koops, K. (2021). The evolutionary origins of primate scleral coloration. bioRxiv, 2021.2007.2025.453695.

18. Yorzinski, J.L., and Miller, J. (2020). Sclera color enhances gaze perception in humans. PLOS ONE 15, e0228275.

19. Tomonaga, M., and Imura, T. (2010). Visual search for human gaze direction by a chimpanzee (*Pan troglodytes*). PLOS ONE 5, e9131.

20. Ricciardelli, P., Baylis, G., and Driver, J. (2000). The positive and negative of human expertise in gaze perception. Cognition 77, B1–B14.

21. Povinelli, D.J., and Eddy, T.J. (1996). Joint visual attention. Psychol. Sci. 7, 129–135.

22. Tomonaga, M. (2007). Is chimpanzee (Pan troglodytes) spatial attention reflexively triggered by the gaze cue? J. Comp. Psychol. 121, 156–170.

23. Endler, J.A. (2008). On the measurement and classification of colour in studies of animal colour patterns. Biological Journal of the Linnean Society 41, 315–352.

24. Egger, H.L., Pine, D.S., Nelson, E., Leibenluft, E., Ernst, M., Towbin, K.E., and Angold, A. (2011). The NIMH Child Emotional Faces Picture Set (NIMH-ChEFS): a new set of children’s facial emotion stimuli. International journal of methods in psychiatric research 20, 145–156.

25. Kaplan, G., and Rogers, L.J. (2002). Patterns of gazing in orangutans (*Pongo pygmaeus*). Int. J. Primatol. 23, 501–526.

26. Farroni, T., Massaccesi, S., Pividori, D., and Johnson, M.H. (2004). Gaze following in newborns. Infancy 5, 39–60.

27. Brooks, R., and Meltzoff, A.N. (2002). The importance of eyes: how infants interpret adult looking behavior. Dev. Psychol. 38, 958–966.

28. Ross, C.F., and Kirk, E.C. (2007). Evolution of eye size and shape in primates. J. Hum. Evol. 52, 294–313.

29. Bekerman, I., Gottlieb, P., and Vaiman, M. (2014). Variations in eyeball diameters of the healthy adults. Journal of Ophthalmology 2014, 503645.

30. Kirk, E.C. (2004). Comparative morphology of the eye in primates. The Anatomical Record Part A: Discoveries in Molecular, Cellular, and Evolutionary Biology 281, 1095–1103.

31. Barr, D.J., Levy, R., Scheepers, C., and Tily, H.J. (2013). Random effects structure for confirmatory hypothesis testing: Keep it maximal. Journal of Memory and Language 68, 255–278.

