## Supplemental Tables and Figures for "Experimental evidence for the gaze-signaling hypothesis: White sclera enhances the visibility of eye-gaze direction in humans and chimpanzees"

5 Supplemental Tables and 2 Supplemental Figures

Table S1. The results from GLMM in Study 1 and 2.

| Experiment | Participant | Effect | $\beta$ | $SE$ | $\chi^2$ | $d.f.$ | $p$ | $\alpha^{*1}$ |
|---|---|---|---|---|---|---|---|---|
| Study 1 Exp. 1 | Humans | Level × Species[*2] | 0.29 (L1 vs. L2) | 1.96 (L1 vs. L2) | | | | |
| | | | 1.09 (L1 vs. L3) | 1.88 (L1 vs. L3) | 3.47 | 3 | 0.32 | 0.05 |
| | | | -0.17 (L1 vs. L4) | 1.76 (L1 vs. L4) | | | | |
| | | Level | -1.04 (L1 vs. L2) | 0.52 (L1 vs. L2) | | | | |
| | | | -2.43 (L1 vs. L3) | 0.50 (L1 vs. L3) | 45.85 | 3 | $< 10^{-9}$ | 0.05 |
| | | | -3.76 (L1 vs. L4) | 0.56 (L1 vs. L4) | | | | |
| | | Species | 2.83 | 0.35 | 37.08 | 1 | $< 10^{-8}$ | 0.05 |
| Study 1 Exp. 2 | Humans | Level × Species × Color | 3.35 | 0.88 | 17.57 | 1 | $< 10^{-4}$ | 0.05 |
| Post-hoc (L3, normal) | | Species | 3.47 | 0.69 | 40.00 | 1 | $< 10^{-9}$ | 0.05 |
| Post-hoc (L3, inverted) | | Species | 3.02 | 0.65 | 22.28 | 1 | $< 10^{-5}$ | 0.05 |
| Post-hoc (L4, normal) | | Species | 1.05 | 0.29 | 30.17 | 1 | $< 10^{-7}$ | 0.05 |
| Post-hoc (L4, inverted) | | Species | 0.47 | 0.27 | 2.91 | 1 | 0.088 | 0.05 |
| Post-hoc (human, L3) | | Color | 3.82 | 0.51 | 30.77 | 1 | $< 10^{-7}$ | 0.05 |
| Post-hoc (chimp, L3) | | Color | 3.27 | 0.71 | 19.74 | 1 | $< 10^{-5}$ | 0.05 |
| Post-hoc (human, L4) | | Color | 1.94 | 0.35 | 17.86 | 1 | $< 10^{-4}$ | 0.05 |
| Post-hoc (chimp, L4) | | Color | 0.92 | 0.25 | 7.89 | 1 | 0.005 | 0.05 |
| Study 2 Exp. 1 (Test A1) | Natsuki | Species | 0.55 | 0.16 | 8.28 | 1 | 0.004 | 0.05/3 |
| | Hatsuka | Species | 0.71 | 0.19 | 9.50 | 1 | 0.002 | 0.05/3 |
| | Pendesa | Species | 1.69 | 0.24 | 21.94 | 1 | $< 10^{-5}$ | 0.05/3 |
| Study 1 Exp. 1-2 (Test A1 vs. B) | Natsuki | Phase × Species | 3.40 | 0.41 | 34.61 | 1 | $< 10^{-8}$ | 0.05/2 |

| | | | | | | | | |
|---|---|---|---|---|---|---|---|---|
| | Hatsuka | Phase × Species | 1.62 | 0.40 | 8.39 | 1 | 0.004 | 0.05/2 |
| Study 1 Exp. 2 (Test B vs. A2) | Natsuki | Phase × Species | 8.49 | 0.86 | 37.04 | 1 | $< 10^{-8}$ | 0.05/2 |
| | Hatsuka | Phase × Species | 3.42 | 0.75 | 33.75 | 1 | $< 10^{-8}$ | 0.05/2 |
| Post-hoc (Test B) | Natsuki | Species | 2.30 | 0.48 | 12.83 | 1 | 0.003 | 0.05/2 |
| | Hatsuka | Species | 0.13 | 0.40 | 0.10 | 1 | 0.75 | 0.05/2 |
| Post-hoc (Test A2) | Natsuki | Species | 0.92 | 0.30 | 7.24 | 1 | 0.007 | 0.05/2 |
| | Hatsuka | Species | 1.46 | 0.27 | 15.11 | 1 | 0.0001 | 0.05/2 |
| Post-hoc (human, Test A1 vs. B) | Natsuki | Species | 0.68 | 0.24 | 7.03 | 1 | 0.008 | 0.05/2 |
| | Hatsuka | Species | 0.09 | 0.21 | 0.18 | 1 | 0.67 | 0.05/2 |
| Post-hoc (human, Test B vs. A2) | Natsuki | Species | 1.25 | 0.42 | 5.33 | 1 | 0.021 | 0.05/2 |
| | Hatsuka | Species | 0.60 | 0.22 | 7.22 | 1 | 0.007 | 0.05/2 |
| Post-hoc (chimp, Test A1 vs. B) | Natsuki | Species | 2.15 | 0.29 | 24.74 | 1 | $< 10^{-6}$ | 0.05/2 |
| | Hatsuka | Species | 0.85 | 0.26 | 9.57 | 1 | 0.002 | 0.05/2 |
| Post-hoc (chimp, Test B vs. A2) | Natsuki | Species | 1.99 | 0.002 | 33.22 | 1 | $< 10^{-8}$ | 0.05/2 |
| | Hatsuka | Species | 1.09 | 0.27 | 14.89 | 1 | 0.0001 | 0.05/2 |

*1 Alpha level was adjusted for the number of individuals in Study 2.

*2 These nonsignificant interaction terms were dropped to test the main effects in these models.

Table S2. Details about the chimpanzee participants.

| Participant | Group | Sex | Age | Rearing condition | Participated in |
|---|---|---|---|---|---|
| Ai | PRI | F | 41 | Nursery/Peers*1 | Training |
| Ayumu | PRI | M | 20 | Mother*1 | Training |
| Chloe*2 | PRI | F | 40 | Nursery/Peers | Training |
| Cleo | PRI | F | 20 | Mother | Training |
| Pal | PRI | F | 20 | Mother | Training |
| Pendesa | PRI | F | 43 | Nursery/Peers | Training, Experiment 1 |
| Hatsuka | KS | F | 12 | Nursery/Peers | Training, Experiment 1-2 |
| Iroha | KS | F | 12 | Mother | Training |
| Mizuki | KS | F | 24 | Nursery/Peers | Training |
| Natsuki | KS | F | 15 | Mother | Training, Experiment 1-2 |

*1 Nursery/Peers indicates the individuals reared by human caretakers and peer conspecifics, while Mother indicates those reared by their biological mothers

*2 Chloe was involved in a related gaze-direction search task in a previous study [1].

*3 Two additional chimpanzees (Zamba and Misaki) participated in a few pilot sessions but did not participate in the training sessions due to low motivation.

Table S3. Details about each training and test stage for the chimpanzees.

| Training/Test phase | Training/Test stage | Description | Number of stimulus individual | Stimulus properties | |
|---|---|---|---|---|---|
| | | | | Size (width × height in pixel) | Brightness (0-100% of the original RGB values) |
| | Training 1 | Presenting stimuli in which the iris was removed (i.e. only sclera was visible in the eye). | | | |
| | Training 2 | Presenting stimuli in which eyes were averted 38 degrees (the iris was visible in the corner of the eye). | 4 (2 chimpanzees, 2 humans) | | |
| Training | Training 3 | Presenting stimuli in which the eyes were averted 30 degrees. | | | |
| | Training 4 | Presenting stimuli in which the eyes were averted 20 degrees (the final position of the iris). | | 400 × 100 | 100 |
| | Training 5 | Presenting 4 new stimulus individuals in half of the trials and 4 old stimulus individuals in the other half. | 8 (4 chimpanzees, 4 humans) | | |
| | Training 6 | | | | |
| Pre-Test A1/A2 | L1 Normal | Presenting L1 stimuli (original size and brightness) in each session. Eye colors of all stimuli are normal. | | | |
| | L1.5 Normal | Presenting L1 stimuli in 24 baseline trials and stimuli with a higher level (smaller and darker) in 24 test trials. Eye colors of all stimuli are normal. | 12 (6 chimpanzees, 6 humans) | 300 × 75 (in test trial) | 75 (in test trial) |
| | L2 Normal | | | 200 × 50 (in test trial) | 50 (in test trial) |
| | L2.5 Normal | | | 150 × 37.5 (in test trial) | 42 (in test trial) |
| Test-A1/A2 | L3 Normal | | | 100 × 25 (in test trial) | 33 (in test trial) |
| | L3.5 Normal | | | 75 × 18.75 (in test trial) | 29 (in test trial) |

| | | | | |
|---|---|---|---|---|
| | L4 Normal | | 50 × 12.5 (in test trial) | 25 (in test trial) |
| | L1 Inverted | | 400 × 100 | 100 |
| Pre-Test B | L1.5 Inverted | | 300 × 75 (in test trial) | 75 (in test trial) |
| | L2 Inverted | | 200 × 50 (in test trial) | 50 (in test trial) |
| | L2.5 Inverted | Same as L1-4 Normal except that eye colors of all images are inverted. | 150 × 37.5 (in test trial) | 42 (in test trial) |
| | L3 Inverted | | 100 × 25 (in test trial) | 33 (in test trial) |
| Test-B | L3.5 Inverted | | 75 × 18.75 (in test trial) | 29 (in test trial) |
| | L4 Inverted | | 50 × 12.5 (in test trial) | 25 (in test trial) |

Table S4. Number of sessions in each training stage.

| Participant | Stage | Number of sessions |
|---|---|---|
| Ai | Training 1 | 4*1 |
| | Training 2 | 51 |
| | Training 3 | 15 |
| | Training 4 | 28 |
| | **Total** | **98** |
| Ayumu | Training 1 | 23*1 |
| | Training 2 | 7 |
| | **Total** | **30** |
| Chloe | Training 1 | 2 |
| | Training 2 | 42 |
| | **Total** | **44** |
| Cleo | Training 1 | 9 |
| | Training 2 | 21 |
| | **Total** | **30** |
| Pal | Training 1 | 26 |
| | **Total** | **26** |
| Pendesa*2 | Training 1 | 2 |
| | Training 2 | 29 |
| | Training 3 | 8 |
| | Training 4 | 14 |
| | Training 5 | 11 |
| | Training 6 | 50 |
| | **Total** | **114** |
| Hatsuka*2 | Training 1 | 15 |
| | Training 2 | 13 |
| | Training 3 | 7 |
| | Training 4 | 11 |
| | Training 5 | 5 |
| | Training 6 | 3 |
| | **Total** | **54** |
| Iroha | Training 1 | 33 |
| | Training 2 | 46 |
| | Training 3 | 12 |
| | **Total** | **91** |

| | | |
|---|---|---|
| Mizuki | Training 1 | 14 |
| | Training 2 | 98 |
| | **Total** | **112** |
| Natsuki[*2] | Training 1 | 4 |
| | Training 2 | 26 |
| | Training 3 | 5 |
| | Training 4 | 15 |
| | Training 5 | 5 |
| | Training 6 | 2 |
| | **Total** | **57** |

*1 Ai and Ayumu were mistakenly moved to Training 2 after only one session scoring >80% in both chimpanzee and human trials (i.e. one additional session was necessary to pass the criteria). For Ai, we performed one additional Training 1 session during Training 2, confirmed that she scored >80% in both chimpanzee and human trials, and then continued her training. Ayumu performed 7 Training 2 sessions after Training 1, but due to his low motivation to participate in this experiment, we decided to drop him from further tests (we also dropped those 7 Training 2 sessions from the analysis).

*2 These three individuals passed all the training stages.

Table S5. Number of sessions in each pre-test and test stage.

| Participant | Test phase | Stage | Number of session |
|---|---|---|---|
| Pendesa | Pre-Test A1 | L1 Normal | 12 |
| | | L1.5 Normal | 4 |
| | | L2 Normal | 19 |
| | | **Total** | **35** |
| | Test-A1 | L2.5 Normal | 26 |
| | | **Total** | **26** |
| Hatsuka | Pre-Test A1 | L1 Normal | 5 |
| | | L1.5 Normal | 8 |
| | | L2 Normal | 3 |
| | | **Total** | **16** |
| | Test-A1 | L2.5 Normal | 13 |
| | | L3 Normal | 13 |
| | | **Total** | **26** |
| | Pre-Test B | L1 Inverted | 10 |
| | | L1.5 Inverted | 2 |
| | | L2 Inverted | 4 |
| | | **Total** | **16** |
| | Test-B | L2.5 Inverted | 2 |
| | | L3 Inverted | 24 |
| | | **Total** | **26** |
| | Pre-Test A2 | L2 Normal | 1 |
| | | L2.5 Normal | 1[*1] |
| | | **Total** | **2** |
| | Test-A2 | L3 Normal | 3 |
| | | L3.5 Normal | 17 |
| | | **Total** | **20** |
| Natsuki | Pre-Test A1 | L1 Normal | 4 |
| | | L1.5 Normal | 5 |
| | | L2 Normal | 6 |
| | | L3 Normal | 2[*2] |
| | | **Total** | **17** |
| | Test-A1 | L2.5 Normal | 15 |
| | | L3 Normal | 18 |
| | | **Total** | **33** |

| | | |
|---|---|---|
| Pre-Test B | L1 Inverted | 10 |
| | L1.5 Inverted | 10 |
| | L2 Inverted | 3 |
| | **Total** | **23** |
| Test-B | L2.5 Inverted | 4 |
| | L3 Inverted | 20 |
| | **Total** | **24** |
| Pre-Test A2 | L1.5 Normal | 1 |
| | L2 Normal | 1 |
| | L2.5 Normal | 1[*1] |
| | **Total** | **3** |
| Test-A2 | L3 Normal | 8 |
| | L3.5 Normal | 18 |
| | L4 Normal | 6 |
| | **Total** | **32** |

*1 These L2.5 sessions in the Test-A2 phase were performed to confirm that the participants' performances did not drop significantly from those in the Test-A1 phase. Test-A2 phase started from L3, the level to which those participants reached in the Test-A1 phase.

*2 As Natsuki performed poorly on these first two L3 sessions, we leveled down the stimuli to L2.5. These initial L3 sessions were not included in the analysis (yet including or not including these two sessions did not change the results).

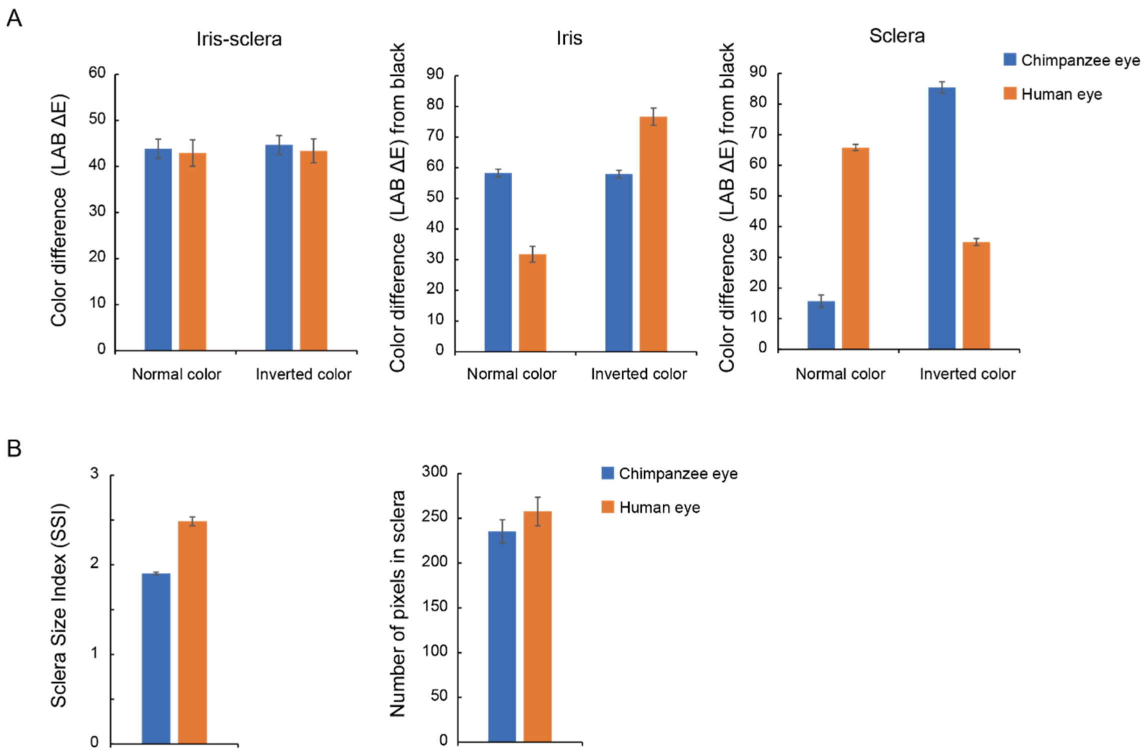


Figure S1. Quantification of stimulus color (A) and shape (B). Color difference was quantified as the Euclidean distance between the two Lab colors $\sqrt{(L1-L2)^2+(a1-a2)^2+(b1-b2)^2}$, following a previous study [2]. The colors of iris and sclera were the means of Lab colors of all pixels respectively in the iris and sclera Regions-Of-Interest (ROI). The iris-sclera color difference was the difference between those two means. The color brightness of iris and sclera was the difference between the mean of each color and the black ($L$ = 0, $a$ = 0, $b$ = 0). The iris-sclera color difference did not significantly differ between the stimulus species in either normal or inverted color ($t$-test; normal: $t_{18}$ = 0.26, $p$ = 0.80, $d$ = 0.12; inverted: $t_{18}$ = 0.39, $p$ = 0.70, $d$ = 0.17). The eye shape was evaluated using Sclera Size Index (SSI), calculated as the longest length of eye opening divided by the iris diameter [3]. The human eye was horizontally longer than the chimpanzee eye, as indicated by higher SSI ($t_{18}$ = 11.34, $p < 10^{-3}$, $d$ = 5.07). We also measured the area size of sclera in the human and chimpanzee eye as the number of pixels in the sclera ROI. The area size did not significantly differ between the stimulus species ($t_{18}$ = 1.08, $p$ = 0.29, $d$ = 0.48). These statistical comparisons were performed using the full stimulus set used for Study 1 (20 images), but the same results were obtained using the stimulus set used for Study 2 (12 images).

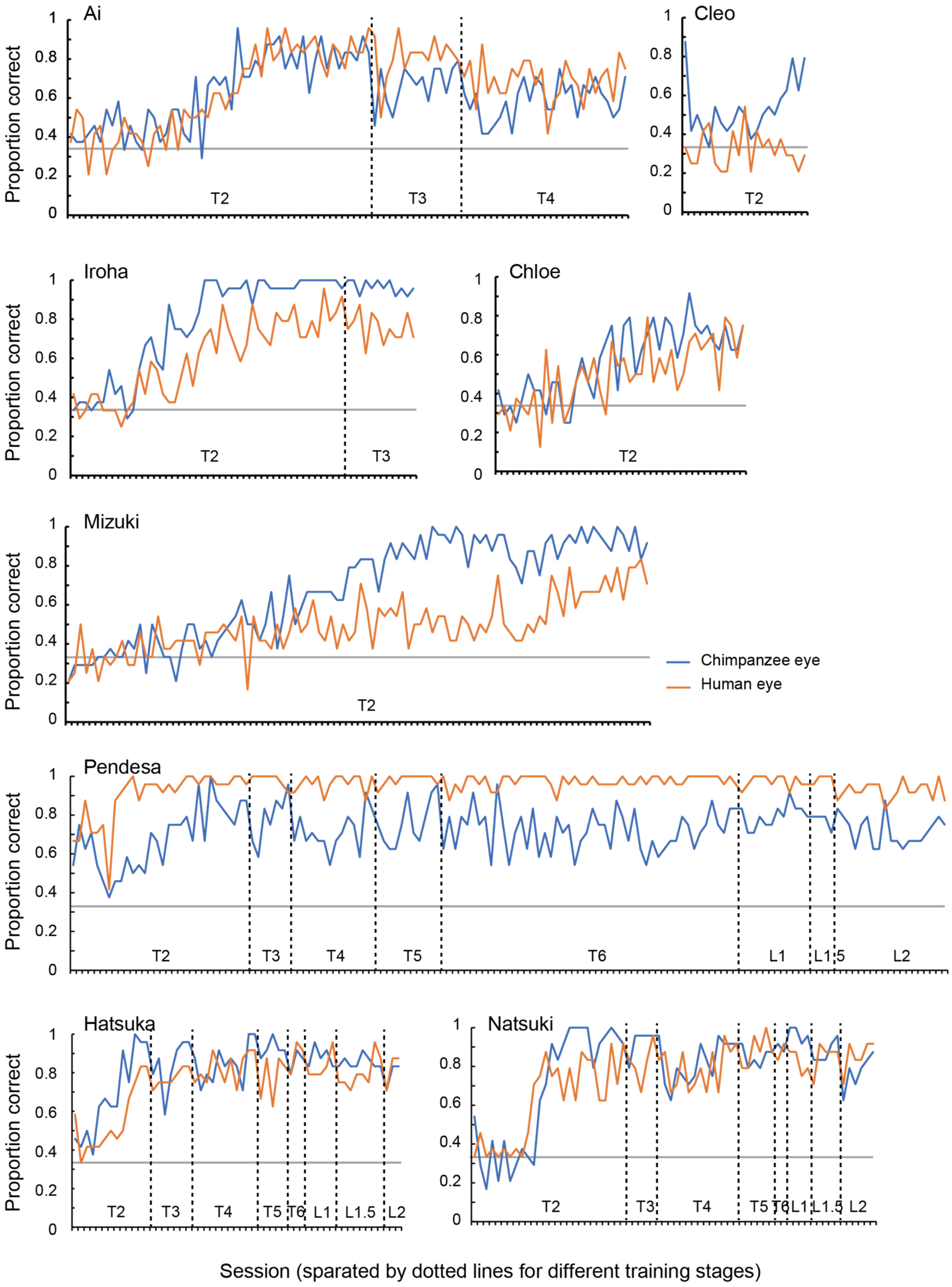


Figure S2. Performance of chimpanzee participants (Ai, Cleo, Chloe, Iroha, Mizuki, Natsuki, Hatsuka, and Pendesa) during Training 2-6 (Training 1 not included because it trained chimpanzees for a visual search task with the no-iris target stimuli, see Table S3) and pre-Test A1 sessions, represented as raw proportion correct across sessions.

References in this Supplemental Material


1. Tomonaga, M., and Imura, T. (2010). Visual search for human gaze direction by a chimpanzee (*Pan troglodytes*). PLOS ONE *5*, e9131.
2. Kano, F., Furuichi, T., Hashimoto, C., Krupenye, C., Leinwand, J.G., Hopper, L.M., Martin, C.F., Otsuka, R., and Tajima, T. (under review, available on request). What is so unique about the human eye? Comparative morphometric, color, and image analysis on the external eye morphology of human and nonhuman great apes and 90 other primate species.
3. Kobayashi, H., and Kohshima, S. (2001). Unique morphology of the human eye and its adaptive meaning: Comparative studies on external morphology of the primate eye. J. Hum. Evol. *40*, 419-435.
